# Predicting trait regulators by identifying co-localization of DNA binding and GWAS variants in regulatory regions

**DOI:** 10.1101/467852

**Authors:** Gerald Quon, Soheil Feizi, Daniel Marbach, Melina Claussnitzer, Manolis Kellis

## Abstract

Genomic regions associated with complex traits and diseases are primarily located in non-coding regions of the genome and have unknown mechanism of action. A critical step to understanding the genetics of complex traits is to fine-map each associated locus; that is, to find the causal variant(s) that underlie genetic associations with a trait. Fine-mapping approaches are currently focused on identifying genomic annotations, such as transcription factor binding sites, which are enriched in direct overlap with candidate causal variants. We introduce CONVERGE, the first computational tool to search for co-localization of GWAS causal variants with transcription factor binding sites in the same regulatory regions, without requiring direct overlap. As a proof of principle, we demonstrate that CONVERGE is able to identify five novel regulators of type 2 diabetes which subsequently validated in knockdown experiments in pancreatic beta cells, while existing fine-mapping methods were unable to find any statistically significant regulators. CONVERGE also recovers more established regulators for total cholesterol compared to other fine-mapping methods. CONVERGE is therefore unique and complementary to existing fine-mapping methods and is useful for exploring the regulatory architecture of complex traits.

## Introduction

Genome-wide association studies (GWAS) have revealed thousands of genetic regions in the human genome associated with diverse complex traits and diseases^1^, but the mechanistic basis of these associations remains largely uncharacterized. With more than 88% of GWAS loci lacking common protein-altering variants, it is increasingly recognized that most GWAS variants may play gene-regulatory roles. Indeed, GWAS variants are enriched in tissue-specific enhancers, thus enabling the prediction of trait-relevant cell types^2–5^, putative causal variants^6–8^ and additional candidate loci^2,9–11^. In particular, a number of methods have been recently developed to exploit regulatory genomics data to fine-map GWAS SNPs^12–14^. One of the principal utilities of these methods is to identify functional annotations, such as transcription factor (TF) binding sites or epigenetic marks, which systematically overlap candidate causal variants.

Modification of transcription factor binding sites of GWAS trait regulators is one proposed mechanism through which non-coding variants overlapping regulatory regions may act. In type 2 diabetes for example, transcription factor binding has been verified to be altered by potential causal variants^15^: TR4 in the ANK1 locus^16^, PRRX1 in the PPARG locus^17^, NEUROD1 at the MTNR1B locus^18^, FOXA1/2 binding at the CAMK1D locus^19^, and ARID5B binding at FTO^20^. However, only FOXA2^18^ and RFX^16^ have been reported as being enriched genome-wide for overlap with T2D variants.

The lack of more globally altered transcription factor binding across GWAS loci led us to hypothesize that some causal variants at GWAS loci may be modifying binding sites of co-factors that in turn interact with trait regulators at the same locus. Under this hypothesis, we expect that causal variants **co-localize** with the binding sites of important regulators in the same regulatory region, but not necessarily overlap them (as is assumed by existing fine-mapping methods). Our hypothesis is consistent with previous observations in fine-mapping of autoimmune diseases, where candidate causal variants were found to have strong tendency to occur proximal to known TF motifs, but not necessarily directly overlap^21^.

We therefore introduce CONVERGE (COmplex trait Networks for Variant, Enhancer and ReGulator Elucidation), a statistical model that identifies significant co-localization of causal variants and TF binding sites within the same regulatory region, while relaxing the strict assumption that causal variants systematically overlap TF binding sites. As a result, CONVERGE is able to identify five novel regulators of T2D that validated in followup experiments, whereas other tested fine mapping methods (fgwas^13^ and LD Score^12^) did not find any enriched TFBS. Furthermore, CONVERGE recovered a similar proportion of established total cholesterol regulators as LD Score, but predicted more established regulators in total. Our trait regulators in turn prioritize novel loci not discovered in the training data but were subsequently identified as genome-wide significant in future and related GWAS studies, and furthermore prioritize suggestive loci more highly than expected by chance. Our trait and tissue-specific predictions of GWAS regulatory regions and upstream trait regulators form important starting points for the systematic experimental dissection of human disease.

## Materials and Methods

### Overview of genomic networks

Our cell type specific genomic networks consist of nodes that represent transcriptional regulators, regulatory regions (enhancers and promoters) and single nucleotide variants. Edges in these networks lead from regulators to regulatory regions and from single nucleotide variants to regulatory regions they tag (either by direct overlap in genomic coordinates or through linkage disequilibrium of neighboring variants). Both node and edge construction are discussed below; unless otherwise specified, each genomic network is constructed independently of all others.

### Construction of network nodes representing regulatory regions, regulators, and single nucleotide variants

From the Roadmap Epigenomics Project^22^, we obtained the locations of enhancers and promoters for each of 127 cell types, that were predicted based on a 15-state ChromHMM model segmentation of histone marks (primarily H3K4me3, H3K4me1 and H3K36me3). For each network, one node is constructed per regulatory region. 67% of regulatory regions show enhancer-associated signatures, compared to 33% that show promoter-associated signatures. One regulator node is constructed for each of 659 regulators for which we could obtain at least one position weight matrix^23^. Finally, one node is constructed for each genetic variant reported in the European population of the 1000 Genomes Project.

### Construction of network edges from regulators to regulatory regions

To identify transcriptional regulators likely to bind each regulatory region, we obtained a set of 1,682 position weight matrices (PWM) spanning 659 transcriptional regulators^23^. Up to 10 shuffled control PWMs were generated for each of the 1,682 true PWMs to estimate the background number of PWM matches expected due to sequence composition alone. For each regulatory region in a given cell type, we scanned the region for matches to each true PWM, aggregating all matches across the entire region to obtain a single total score. We then compute a background (expected) score by averaging aggregate scores computed individually for each of the shuffled versions of that PWM. If the difference between the aggregate score of the true PWM versus the average aggregate score of the shuffled PWMs is positive, the PWM is considered to match the regulatory region with weight equal to the (positive) difference. For each regulator, we then sum the positive difference-scores of the true PWMs belonging to that regulator to estimate a final regulator-regulatory region pair score. We construct network edges from regulators to regulatory regions when this pair score (between regulator and regulatory region) is at least 0.5. Finally, given the *T* × *R* matrix of edges *E* between *T* regulators and *R* regulatory regions, we re-scale all edges to correct weights by degree by setting
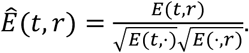
Rescaling helps prevent regulators with high out-degree and regulatory regions with high in-degree (network hubs) from being prioritized simply due to high overall connectivity. Previous work using the same motifs and epigenomic data show that motif enrichments in regulatory regions recapitulates known associations^22^.

### Construction of network edges from single nucleotide variants to tagged regulatory regions

We constructed an initial map of 1000 Genomes variants to regulatory regions by finding all variants directly overlapping regulatory regions in each cell type. Then, using the 1000 Genomes Phase One release 3 European population data^24^, we computed R^2^ values between all pairs of variants (within 1mb of each other) present in the European population. Additional links were then constructed between variants and regulatory regions overlapping SNPs in high LD (R^2^ > 0.8). For each variant in each cell type, the edges leading from that variant to tagged regulatory regions are assigned equal weights that sum to one. Finally, all variant chromosome coordinates were updated to hg19 using dbSNP version 138.

### Validation of regulator-enhancer edges in the network

We downloaded all ChIP-Seq genome-wide profiles for Tier 1+2 cell types from ENCODE on January 10, 2017. We removed those ChIP-Seq profiles corresponding to eGFP conjugated factors, as well as the MCF7 and Sknsh cell lines, as they were not included in the Roadmap Epigenomics Consortium epigenomes. This left eight cell types with ChIP-Seq data to validate our networks (H1, IMR90, A549, GM12878, HELA, HEPG2, HUVEC, K562). After mapping factor names to gene symbols, we only kept those TF ChIP-Seq experiments where the TF also has a corresponding node in the networks. For each TF ChIP-Seq profile, we then used the hypergeometric test to measure the significance of the overlap between ChIP-Seq peaks in enhancers, and the set of enhancer nodes that a given regulator node is connected to. For each TF ChIP-Seq experiment, we record the rank of the matched regulator node in the network relative to all other regulator nodes in the network, and compute the empirical cumulative distribution function to assess and visualize the overall concordance between regulator edges and ChIP-Seq experiments.

### Clustering regulators in each genomic network into regulatory modules

To cluster regulators into groups that bind similar regulatory regions, for each GN we computed a linear dot product kernel *K* of the scaled network matrices *N*, defined as *K* = *N* × *N^T^*. *K* was then used as input to the affinity propagation clustering algorithm^25^, and clustered in a two step procedure. First, it was run with similarity matrix set to *K* and parameters *p* = *q* = NA. After the first round of clustering, we observed that only TFs with similar binding sites group together, though we expect there to exist sets of TFs with similar target regulatory regions despite having different sequence binding preferences. We therefore ran affinity propagation a second time to discover “meta-clusters”, only using the exemplar regulators discovered in the first iteration (and the corresponding submatrix of *K*), again with *p* = *q* = NA. Each regulator was assigned to a meta-cluster based on its exemplar identified in the first iteration of affinity propagation, and that exemplar’s subsequent assignment to a meta-cluster in the second iteration of affinity propagation.

### Genome-wide association P-values

We collected a compendium of meta-GWAS summary statistics for analysis, with particular focus on type 2 diabetes^26^ and lipid traits (HDL, LDL, and total cholesterol)^27,28^. See Supplementary Note 1 for a full list of GWAS. We collected P-value summary statistics (median of 2,462,778 SNPs per study) from their respective web sites, and updated SNP locations and identifiers to hg19 using dbSNP version 138. Only SNPs that were identified in the 1000 Genomes Phase One release 3 were kept. SNPs within 2Mb of the coding region of all HLA genes were removed to identify signal outside of the polymorphic HLA region.

### LD pruning and SNP classification by association P-value

The remaining SNPs from each GWAS (median of 2,451,650) were then pruned by iteratively selecting the SNP with the smallest P-value, then removing all other SNPs within a 1mb window whose pairwise LD (as estimated from the 1000 Genomes Phase One release 3 European population^24^) was greater than 0.2, leaving a median of 40 SNPs per study. We assigned each SNP to one category of significance, distinguishing: (a) genome-wide significant loci, defined as those pruned SNPs whose association P-value is less than 5×10^-8^; (b) suggestive loci, defined as those whose P-value is less than 5×10^-6^, but greater than 5×10^-8^. Finally, for each cell type analysis separately, we merged the pruned GWAS variants whose tagged regulatory regions overlap (by at least one region) into a single agglomerated GWAS variant, such that any regulatory region in the cell type is tagged by zero or one GWAS variants.

### Predicting GWAS trait tissues based on regulatory enrichment in GWAS SNPs

This prediction involves three steps.

#### a) Classification of SNPs into SNP-groups based on genomic features

We generated SNP-groups that group SNPs based on similar genomic features as follows. We first ranked all GWAS variants by (i) minor allele frequency (MAF) in the 1000 Genomes European population; (ii) distance to nearest protein coding gene transcription start site; and (iii) approximate number of SNPs in LD (computed as the sum of all R^2^ values between the query SNP and all other 1000 Genomes SNPs within a distance of 250kb). For each of these three features, the ranked set of variants were divided into 15 approximately equally sized bins, then each SNP was grouped with other SNPs that fall into the same bin for all three features (yielding a total of 15^3^=3,375 SNP-groups).

For each SNP-group, we defined the number of independent loci within that group by: (a) identifying all pairs of SNPs within the SNP-group in linkage disequilibrium (R^2^ >= 0.8); (b) using these pairwise links as edges in a graph, where each node is a member SNP of the SNP-group; and (c) computing the number of connected components in this graph as a conservative estimate of the number of independent loci in that SNP-group (not all connected SNPs within each group are directly in linkage disequilibrium).

#### b) Removal of annotations of SNPs tagging constitutively active regulatory regions

For GWAS tissue-specific enrichment testing below, we identified the set of SNPs that tag constitutive regulatory regions for exclusion because they are uninformative for identifying enrichment of GWAS SNPs in cell type specific regulatory regions. First, we identified all SNPs that tag any regulatory region in each cell type. The 127 cell types have previously been categorized based on tissue of origin^22^, so we defined a SNP as tagging a constitutive regulatory region if it covers at least 75% of the tissue groups, where a tissue group is considered covered if the SNP tags a regulatory region in at least 25% of the member cell types of that tissue group. Specifically in our method for identifying cell type specific enrichment of GWAS SNPs, these constitutively-tagging SNPs are not considered to tag any regulatory region. This resulted in the removal of 9% of SNPs that tag at least one regulatory region in one cell type.

#### c) Permutation testing

For each GWAS trait and each cell type, we use the SNP-groups to count the number of independent suggestive GWAS SNPs (P < 5×10^−6^) that tag a regulatory region in that cell type. We then used our 3,375 SNP-groups to generate permuted SNP sets of the same number of independent loci (and furthermore with the same number of variants from each SNP-group). Our test statistic is the number of independent loci that tag a regulatory region, and we generate 10^6^ permutation sets to compute an empirical P-value for the enrichment of a set of GWAS variants to locate near regulatory regions of a particular cell type. Trait tissues were defined as those GWAS-cell type enrichments passing a per-trait Bonferroni significance cutoff of 0.01.

### Identifying trait-regulators and regulatory modules

Our statistical model CONVERGE is applied independently for each GWAS-trait tissue combination, to infer a trait pathway that consists of a set of trait regulators and trait regulatory regions (a subset of which are targeted by GWAS variants).

CONVERGE assumes the following input data is available and known prior to analysis:

- A trait tissue of the GWAS variants, for which we have already inferred a genomic network *N*. This network *N* connects a set of *T* regulators with a set of *R* regulatory regions, with edge weights *N*_*f*,*r*_ connecting each regulator *f* to each regulatory region *r*, which is inferred as described above.
- A SNP-to-region network *𝑣* where *𝑣*_*s*,*r*_ = 1 if regulatory region *r* is tagged by GWAS variant *s*, else *𝑣*_*s*,*r*_ = 0. From *𝑣*, we can compute a partitioning of the set of all *R* regulatory regions in the network, Ω = {1,…, *R*}, into three distinct groups, e.g., Ω = Ω_bg_ ∪ Ω_fg_ ∪ Ω_unk_: (1) Ω_bg_ represents those ‘background’ regulatory regions that are not tagged by a GWAS variant are assumed to not be in the trait pathway during training; (2) Ω_fg_ represents those ‘foreground’ regulatory regions tagged by a GWAS variant in a 1-1 relationship (e.g. the corresponding GWAS variant only tags a single regulatory region), and therefore are assumed to be the target regions of GWAS variants; and (3) Ω_unk_ represents those ambiguous regulatory regions located in a GWAS locus with multiple other regulatory regions, for which it is unclear which regulatory region in the locus is the target.

From Ω, CONVERGE then defines a set of *R* ‘trait-regulatory region’ variables *θ_r_* indicating whether a given regulatory region r is a trait-regulatory region (*θ_r_* = 1) or not (*θ_r_* = 0). CONVERGE sets *θ_r_* = 0 for background regulatory regions *r* ∈ Ω_bg_ and *θ_r_* = 1 for foreground regulatory regions *r* ∈ Ω_fg_, therefore treating these variables as observed data. From the observed data, CONVERGE infers the following latent variables and model parameters:

- CONVERGE infers the regulatory region indicator variables *θ_r_* for those regions *r* ∈ Ω_unk_ located in GWAS loci tagging multiple regulatory regions. Here the model assumes exactly one regulatory region in each GWAS locus is a trait-regulatory region, and the remaining are zero, e.g. Σ_*r*_*𝑣*_*s*,*r*_*θ_r_* = 1 for all GWAS loci *s*.
- A set of weights *α* of length *T* indicating the influence of each regulator on the trait pathway. Larger *α* indicates more influence and a higher enrichment of that regulator’s targets in GWAS target regions. Trait regulators are defined as all transcriptional regulators whose weight is positive, i.e., *α_f_* > 0.

We use the expectation maximization algorithm to perform inference and learning.

### Expectation maximization (EM) algorithm for learning CONVERGE

For CONVERGE, the EM algorithm can be thought of as iteratively adjusting weights *α_f_* of the regulators of the genomic network that specify the trait regulators, in order to maximize the flow of weight from the regulators to the GWAS variants, while minimizing flow to non-GWAS target regulatory regions (e.g. the background regulatory regions). **Initialization:** We first randomly assign non-zero weights to a subset of regulators to choose the initial trait regulators, and randomly select a target regulatory region for each GWAS SNP. **E-Step:** The current set of weights on the regulators is propagated through the genomic network to the regulatory regions to form prior probabilities, and the target GWAS regulatory regions *θ_r_* (*r* ∈ Ω_unk_) are updated such that regulatory regions with higher prior probability are more likely to be a GWAS target regulatory region. **M-step:** Given the current set of GWAS target regulatory regions *θ_r_*, we re-weigh the regulators *α_f_* to identify a set of trait regulators that best distinguish GWAS target regulatory regions from all other regulatory regions in that cell type. We iterate the E-step and M-step until convergence. Regulators with positive weights *α_f_* after convergence are labeled as trait regulators. See Supplementary Note 2 for full details of the model learning procedure.

### Permuting the genomic networks for statistical significance testing of trait pathways

In order to assess statistical significance of a trait pathway, we generated 500 permutations of each genomic network as follows. We first ranked all regulatory regions by (a) length; (b) number of 1000 Genomes European SNPs overlapping the regulatory region; (c) average minor allele frequency of SNPs overlapping the regulatory region; and (d) average specificity of the regulatory region, defined as the number of cell types that have an overlapping regulatory region. For each of these four features, the ranked set of variants was divided into 5 approximately equally sized bins (except for the specificity feature, which was divided into 10 bins). Each regulatory region was then grouped with other regulatory regions that fall into the same bin for all four features (yielding a total of 10×5^3^=1,250 region-groups). We then generated 500 randomized versions of each GN by permuting the labels of regulatory regions within the same region-group. Our permutation scheme is designed to maintain the in and out degrees of both the regulators and regulatory regions, to ensure statistically significant trait pathways are not due to GWAS variants tagging regulatory regions of high degree.

### Calculating priorities for regulatory regions using trait-regulators

After CONVERGE is trained using a set of GWAS variants and a genomic network, we computed the values *P*(*θ_r_* = 1|*N*,*α*) for all regulatory regions *r*. We then computed *P*_random_max__(*θ_r_* = 1|*N*,*α*), the maximum value of *P*(*θ_r_* = 1|*N*,*α*) achieved from 500 random permutations of the GN (described above). The priority of a regulatory region is defined as *P*(*θ_r_* = 1|*N*,*α*) − *P*_random_max__ (*θ_r_* = 1|*N*,*α*).

### Partitioning variants into those in the trait pathway and those that are not

For each GWAS trait and trait tissue for which we found a statistically significant trait pathway, we then sought to identify which subset of the GWAS variants contributed to the trait pathway and identification of trait regulators (Fig. 2). Within the context of a trait tissue, we therefore partition GWAS variants into one of three categories:

**Figure 1:**
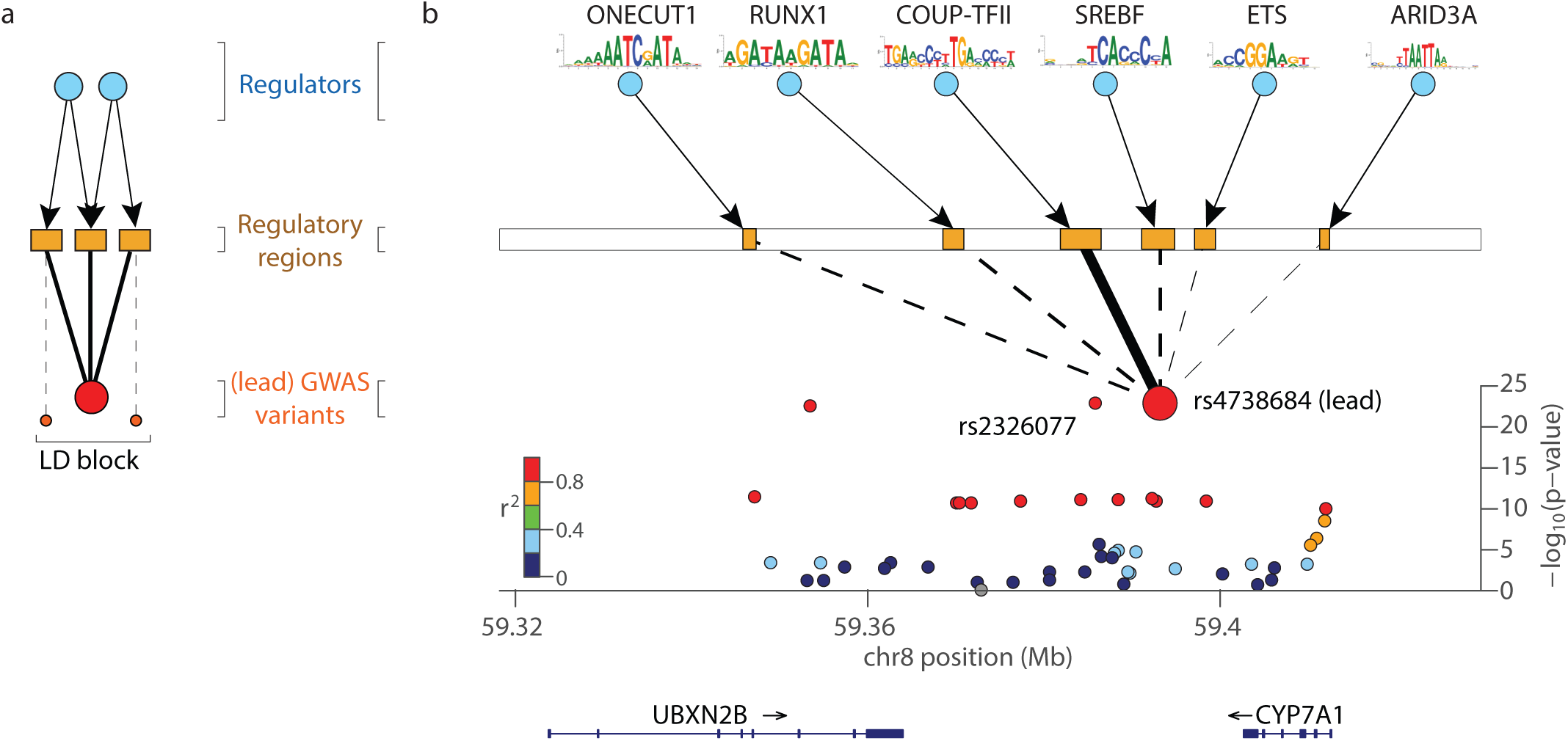
Finding GWAS trait pathways by leveraging genomic networks. **(a) Structure of a genomic network (GN).** GNs are three layer networks consisting of regulators, regulatory regions and GWAS variants. One regulator node is created for each of 659 transcriptional regulators (blue nodes) for which a position weight matrix is available, one variant node is created for each GWAS locus identified by the study (large red nodes), and one regulatory region node is created for every enhancer and promoter (orange rectangles) identified within a cell type by the Roadmap Epigenomics Consortium. Variant nodes are connected to regulatory regions they either directly overlap on the genome, or overlap through another variant (small red circles) by linkage disequilibrium (R^2^>0.8). Regulatory regions are linked to their upstream regulators through motif occurrences within the regulatory region sequence. **(b) The adult liver genomic network centered on the CYP7A1 total cholesterol-associated locus.** Shown is the CYP7A1 locus associated with total cholesterol (lead variant rs4738684, P= 1.12 × 10^−23^). rs4738684 tags six separate enhancers, of which CONVERGE selects the single regulatory region (solid black line) that is bound by the regulator COUP-TFII whose binding sites are also found at other GWAS loci. The predicted target enhancer also overlaps another variant (rs2326077, P=1.12 × 10^−23^) in LD and which demonstrates equally strong association as the lead variant, even though association strength was not used as input to the model. COUP-TFII regulation of CYP7A1 is a previously reported regulatory interaction in the literature.

1. No regulatory activity: variants that do not tag any regulatory region in the trait tissue genomic network are ignored and assumed not to belong to the trait pathway.
2. Regulatory activity at locus, not in trait pathway: variants that tag at least one regulatory region in the trait tissue, but when the network is permuted, the regulatory region is assigned a priority at least as large as its priority under the real network, in at least 5% of the permutations.
3. Regulatory activity at locus, in trait pathway: variants that tag at least one regulatory region in the trait tissue, and when the network is permuted, the regulatory region is assigned a priority at least as large as its priority under the real network, in less than 5% of the permutations.

### Calculating the regulator z-scores and regulator network ranks

Trait regulators are defined as those regulators whose weights *α_f_* are estimated by CONVERGE to be greater than 0. To quantify the extent to which each regulator is specific to the trait (and not simply a hub in the GN), we computed a regulator z-score, defined as the z-score of the regulator weight (computed on the real network) relative to the mean and standard deviation of the weight assigned to the same regulator under 500 permutations of the network (described above). We also compute a regulator network rank, defined as the regulator’s relative rank with respect to the fraction of times it is selected as a trait regulator in the 500 permutations of the corresponding network (described above). A network importance score of 1.0 means the regulator was most often selected in the permuted networks than any other regulator, whereas a score of 0 means the regulator was selected the least frequently in the permuted networks.

### siRNA knockdown of T2D factors and assessment of glucose-stimulated insulin secretion in pancreatic INS-1 β cells

T2D regulators were identified as all regulators whose weights *α_f_* are estimated by CONVERGE to be greater than 0. We identified negative control regulators by permuting the network (in a node-degree sensitive way described earlier) 500 times, averaging the weights *α_f_* learned for each factor across the 500 permutations as described above, then selecting the factors with highest average weight as negative controls. The rat insulinoma cell line INS-1 was cultured in RPMI medium (supplemented with 10% fetal bovine serum, 100 mM sodium pyruvate, penicillin/streptomycin and 50 μM 2-mercaptoethanol). Cells were treated with 25nM nontargeting (NT) control or siRNA targeting the predicted T2D regulators (HOXA9, ZSCAN4, OTX1, EGR, ID4, SMAD4), as well as the negative controls (NEUROD2, ITGB2, ZNF384, GFI1, GATA1, BATF3) (ON-TARGETplus human siRNA SMARTpool (Dharmacon, USA)) using HiPerFect (Qiagen, Germany) according to the manufacturer’s protocol. After 72 hours, the medium was changed to low glucose concentration (5 mM) for 24 h. On the next day the medium was changed to low glucose (5mM) or high glucose medium (25mM) (stimulated) for 1 hour to induce glucose-stimulated insulin-secretion. The medium supernatant was collected and insulin concentrations were measured using a commercially available insulin-ELISA (Mercodia, Sweden). The cells were harvested in buffer RLT (Qiagen, Germany) and frozen at −80 C for extraction of RNA and determination of knockdown efficiency. We focused on a cellular phenotype that is amenable to quantitative evaluation, namely glucose-stimulated insulin secretion capacity of pancreatic insulinoma INS-1 beta cells, measured using the enzyme-linked immunosorbent assay, and used it to evaluate the trait regulators predicted for type 2 diabetes in the pancreatic islet network (Fig. 3b). For siNT, OTX1, ID4, EGR, ZSCAN4, HOXA9 and SMAD4, seven replicates were successful. Five replicates were successful for NEUROD2 and ITGB2, while four replicates were successful for ZNF384, GFI1, GATA1, and BATF3. Increased levels of glucose in the circulation lead to increased glucose uptake into pancreatic beta cells, which in turn leads to increased insulin secretion. This process is impaired type 2 diabetic patients and was therefore selected as the phenotype of interest.

### Predicting novel GWAS loci

For the LDL, HDL, total cholesterol and triglyceride level traits, we were able to obtain GWAS summary statistics collected at two different dates, one in 2010^28^ and one in 2013^27^. Prediction consisted of two steps:

#### a) Training

We used the historical data (2010) to train CONVERGE and prioritize regulatory regions. We identified statistically significant trait pathways in the liver network for HDL, LDL and TC traits, and therefore prioritized all liver regulatory regions with respect to the trait regulators for each of those three traits.

#### b) Evaluation

For each of the three traits, we identified all “novel” regulatory regions tagged by the genome-wide significant loci from the more recent study (2013) but were not tagged by any of the genome-wide significant loci in the historical study (2010). We hypothesized that these novel regulatory regions should be enriched for true GWAS target regulatory regions. We therefore evaluated each of the three traits’ regulatory region prioritizations both by how highly ranked these novel regulatory regions were on average (AUROC), and whether the top ranked regions specifically were enriched in novel regulatory regions (AUPR). To assess prediction performance, we also measured these AUROC and AUPR performance measures when liver regulatory regions were prioritized using the permuted networks.

### Predicting suggestive GWAS loci

We also tested CONVERGE prioritization of regulatory regions for their ability to predict suggestive GWAS loci. Similar to our procedure for testing novel GWAS loci, for each combination of GWAS trait and trait tissue for which a statistically significant trait pathway was identified, we then identified all regulatory regions tagged by suggestive GWAS loci (5 × 10^−8^ < *P* < 5 × 10^−6^) but not tagged by the genome-wide significant GWAS loci. Unlike the prediction of novel loci, where we expect most loci to be real, we reasoned that many regulatory regions tagged by suggestive GWAS loci were not bona fide, and therefore the appropriate accuracy measure in this experiment (AUPR) is one that yields high values when the most prioritized regulatory regions are enriched for some of the suggestive loci, as opposed to a measure that yields high values when all suggestive loci are more highly prioritized overall compared to other regulatory regions (e.g. AUROC).

### Estimating the number of additional trait regulatory regions in the genome

To estimate the total size of the regulatory architecture of each trait, we counted the total number of regulatory regions in each target cell type that were not tagged by GWAS loci but exceeded a priority threshold set by regulatory regions tagged by suggestive GWAS loci. That is, under the assumption that at least one of the suggestive GWAS loci is bona fide (and therefore a regulatory region in its locus is a true trait regulatory region), we used the maximum priority value obtained by any regulatory region tagged by a suggestive GWAS locus as the threshold for identifying novel candidate regulatory regions.

### Transcription factor and cell type enrichment testing with fgwas

Fgwas (version 0.3.6) was run with default parameters and using the suggested workflow in the manual for testing individual annotation enrichment, except in cases where the summary statistics explicitly list the number of cases and controls, in which case the –cc option was used^13^. When the summary statistics released for each GWAS trait did not include the number of cases and controls used for testing each SNP, we used the number of cases and controls reported in the original study. When no Z-score or direction of effect was reported (only a P-value given), we converted the P-value to a Z-score. Because conversion of P-values from summary statistics to Z-scores requires use of the qnorm() function in R, and qnorm returns –Inf for p-values smaller than 1e-323, we approximate the qnorm of smaller pvalues with the qnorm of 1e-323. Minor allele frequency, when not reported in the original summary statistics, were replaced with values computed from the 1000 Genomes Phase One data. SNPs from summary statistics were further restricted to those in the 1kG European population, so all traits were treated homogeneously. Enrichment of a specific annotation for a GWAS trait was computed using a likelihood ratio test, comparing the likelihood of fgwas fit with an annotation, compared with the baseline (no annotation). For identifying enriched cell types, we used the same Roadmap epigenome annotations used for CONVERGE described above. For TF enrichments, we used two sets of transcription factor binding sites. The first set we generated by identifying all transcription factor motif matches across the human genome using the same set of 659 motifs used to generate our genome networks, except we only keep motifs that have a conservation score of at least 0.5^23^ (“Set 1”). The second set of motif instances we use are provided by Kitchaev et al. for 165 factors^14^ (“Set 2”).

### Transcription factor and cell type enrichment testing with LD Score (LD Score)

For each input file of GWAS summary statistics from the original studies, we used the munge_sumstats.py script from LD Score^12^ version 1.0.0 to extract the relevant summary statistics. For identifying enriched cell types, we used the same Roadmap epigenome annotations used for CONVERGE described above. We performed TF enrichment testing while controlling for all 53 categories of cell type-agnostic annotations in the baseline model^29^, as outlined in the LD Score manual, and retrieving the P-values associated with each of our annotations. Transcription factor enrichment testing used the Set 1 and Set 2 TF annotations described above for fgwas.

## Results: Inference of 127 genomic networks

Linkage disequilibrium (LD) is a barrier to identifying co-localized TFBS and GWAS variants, as a GWAS SNP can tag multiple regulatory regions. We therefore construct genomic networks linking regulators, regulatory regions and GWAS SNPs to help identify GWAS target regulatory regions and upstream regulators. Previous studies constructing networks linking regulators to targets focused on using either ChIP-Seq profiles from multiple cell types to construct a single human regulatory network^30^, or constructed cell type-specific TF-TF networks by computationally predicting the binding locations of TFs in the promoter regions of other TFs^31^. Neither of these are applicable to our study because they are either not cell type-specific or do not consider distal regulatory elements where GWAS loci may reside. Therefore, we began by linking GWAS variants into 127 distinct tissue and cell type-specific genomic networks (GNs) (Fig. 1), each consisting of:

- Regulatory region nodes representing tissue-specific regulatory regions^22^, of which there are a median of 146,754 regulatory regions in a given genomic network (Supplementary Fig. 1).
- Variant nodes representing common genetic variants.
- Regulator nodes representing sequence-specific transcription factors (TFs).
- Variant edges connecting genetic variants to the regulatory regions whose activity they may affect. These reflect either direct overlap in genomic coordinates of variants and regulatory regions, or indirect overlap when a variant is in LD with another variant overlapping a regulatory region.
- Regulator edges connecting regulators to the regulatory regions they bind, as predicted by their known DNA sequence preferences. On average, regulatory regions are linked to 8 regulators, and each regulator targets ~1,700 regions (Supplementary Fig. 2).

We validated the inferred regulator-regulatory region edges in our networks by comparing them to all available TF ChIP-Seq genome-wide profiles generated by the ENCODE consortium (Supplementary Fig. 3). For each cell type for which ChIP-Seq data was available and we inferred a genomic network, we assessed the statistical significance of the overlap of each ChIP-Seq profile (corresponding to peaks in one cell type for one TF) and the edges from each individual regulator node in the network. For each ChIP-Seq profile, we used the rank of the correctly matched node in the network to assess the accuracy of the network, where smaller rank indicates a more relatively significant overlap between the correct TF and the corresponding ChIP-Seq profile. Across all tested cell types, the correct regulator node was within the top 5 most significant overlapping TFs for 50% of the ChIP-Seq experiments tested, and was within the top 50 most significant overlapping TFs for 80% of the ChIP-Seq experiments.

Based on network connectivity, GWAS variants are predicted to have diverse impact on gene regulation depending on the particular tissue in which it acts. In the 42 GWAS we analyzed, GWAS variants tagged up to 49 distinct regulatory regions in a given cell type, with a median of 2 regions per regulatory variant (Supplementary Fig. 4). Furthermore, of variants tagging at least one region, 53% tagged more than one regulatory region per tissue type (ambiguous GWAS variants) (Supplementary Fig. 5), highlighting the ambiguity of the mechanism of action of many GWAS variants. Regulatory regions tagged by ambiguous GWAS variants have an average pairwise similarity (Jaccard index) of 3% with respect to their upstream regulators, further demonstrating the diversity of possible functional consequences of GWAS variants.

## Results: Predicted trait tissues of GWAS variants distinguish related phenotypes

To further validate the structure of the genomic networks, we used the SNP-regulatory region links to predict trait-relevant tissues of GWAS, in a manner analogous to previous approaches that identify overlap between GWAS variants and regulatory annotations for individual tissues^12–14,21,22,32^. The regulatory architecture of a complex trait and its associated variants is expected to be dependent on context, for example the trait tissues in which the variants act, and identifying trait tissues allows us to substantially reduce the number of candidate variants for each of these traits by focusing on those overlapping regulatory regions active in trait tissues. Here we designed a test that specifically uses our genomic networks, so that we can validate the correctness of the network structure. Our SNP permutation-based approach uses random sets of GWAS SNPs to control for minor allele frequency, distance to transcriptional start site and LD block size. To identify only the strongest tissue enrichments, we removed the constitutive regulatory region annotations (see Methods), which led to an increase in specificity of our predictions at the cost of power.

We collected a comprehensive catalog of case-control and quantitative GWAS summary statistics (Supplementary Table 1), and predicted the trait tissues for each GWAS (Fig. 2a). We found that the predicted GWAS trait tissues agree well with our current physiological understanding of the etiology of these diseases and traits (Fig. 2a), and were able to recognize important differences in the tissue/cell types underlying related traits known to affect distinct biological processes.

**Figure 2:**
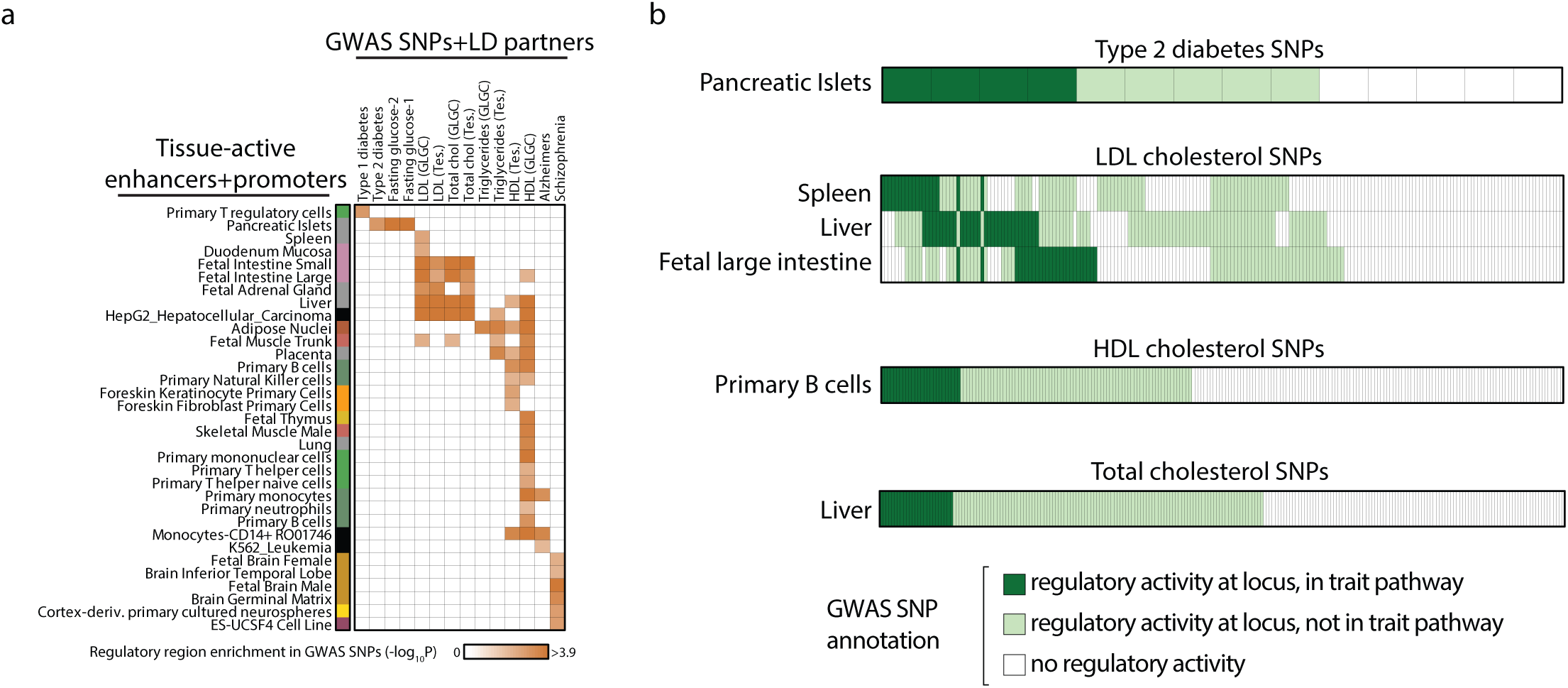
Identification of GWAS trait pathways within targeted cell types. **(a) Identification of trait tissues of GWAS variants.** GWAS trait tissues (row) for each GWAS (column) were identified through permutation testing and are indicated by the orange squares. The orange shading is proportional to the statistical significance (after Bonferroni correction of P-values calculated from permutation testing) of the enrichment. Only P-values smaller than 0.01 are shaded. Cell types (rows) are colored based on their cell type and tissue of origin^22^. (**b**) **GWAS variants target distinct trait pathways in different trait tissues.** For each trait tissue of each GWAS identified in (a), we applied CONVERGE to detect GWAS trait pathways, and used permutation testing to identify statistically significant pathways. Each heatmap indicates a GWAS trait for which at least one significant pathway was found, and each row represents a trait tissue in which a pathway was found, where individual columns represent single GWAS variants. GWAS variants are partitioned into three categories: variants which did not tag any regulatory region in that cell type and therefore are assumed to not participate in the trait pathway in that cell type (white); variants which did tag at least one regulatory region but were not bound with high specificity to the trait regulators and therefore are predicted to not be in the trait pathway (light green); and variants that did tag at least one regulatory region and were bound with high specificity to the trait regulators, and therefore were inferred to be part of the pathway (dark green). For cholesterol traits, only enrichments for the newer study (2013) are shown.

First, we recovered known important differences between low-density lipoprotein and high-density lipoprotein cholesterol (LDL vs. HDL). LDL-associated variants target primary liver and intestine tissue, consistent with both intestinal and hepatic regulation of cholesterol absorption, hepatic lipoprotein remodeling, and LDL cholesterol synthesis after cholesterol intake via bile acid synthesis and excretion^33^. In contrast, HDL variants target adipose nuclei and skeletal muscle, consistent with their roles in peripheral tissues^34,35^, and in diverse types of immune cells, including B and T cells, consistent with roles in innate and adaptive immune response, inflammatory response tuning, antigen presentation in macrophages, and B and T cell activation^36^.

Second, we recovered known differences between type 1 diabetes and type 2 diabetes (T1D vs. T2D). T2D-associated variants were only predicted to target pancreatic islets, consistent with dysregulation of insulin secretion processes that control blood glucose levels^37,38^, and similarly to fasting glucose level-associated variants. In contrast, T1D-associated variants only targeted T regulatory cells, indicating dysregulation of immunity, consistent with the well-established auto-immune basis of T1D^39^.

Third, we recovered known differences between Alzheimer’s disease and schizophrenia variants. We found that AD-associated variants target primary monocyte immune cells, consistent with monocyte-specific eQTL enrichment for AD variants^40^, and a recently-recognized immune and inflammatory basis of AD in mouse and human^41^. In contrast, Schizophrenia-associated variants target multiple brain tissues, including fetal brain tissue, brain germinal matrix, neurospheres, and embryonic stem cells, indicating dysregulation of early brain development.

## Results: Bayesian model for predicting GWAS regulatory trait pathways

Having identified trait tissues for each GWAS, we next developed CONVERGE, a probabilistic graphical model to simultaneously identify the target regulatory region in each GWAS locus, as well as the trait regulators that co-localize with the candidate causal variant (Fig. 3a, Supplementary Fig. 6). Our Bayesian model receives as inputs a set of GWAS SNPs and one genomic network corresponding to a trait tissue. In turn, CONVERGE (1) predicts the trait regulators in that cell/tissue type, (2) prioritizes all regulatory regions in the genome with respect to evidence that they are trait regulatory regions based only on trait regulator binding, and (3) predicts the target regulatory region within each GWAS locus (Fig. 1b).

**Figure 3:**
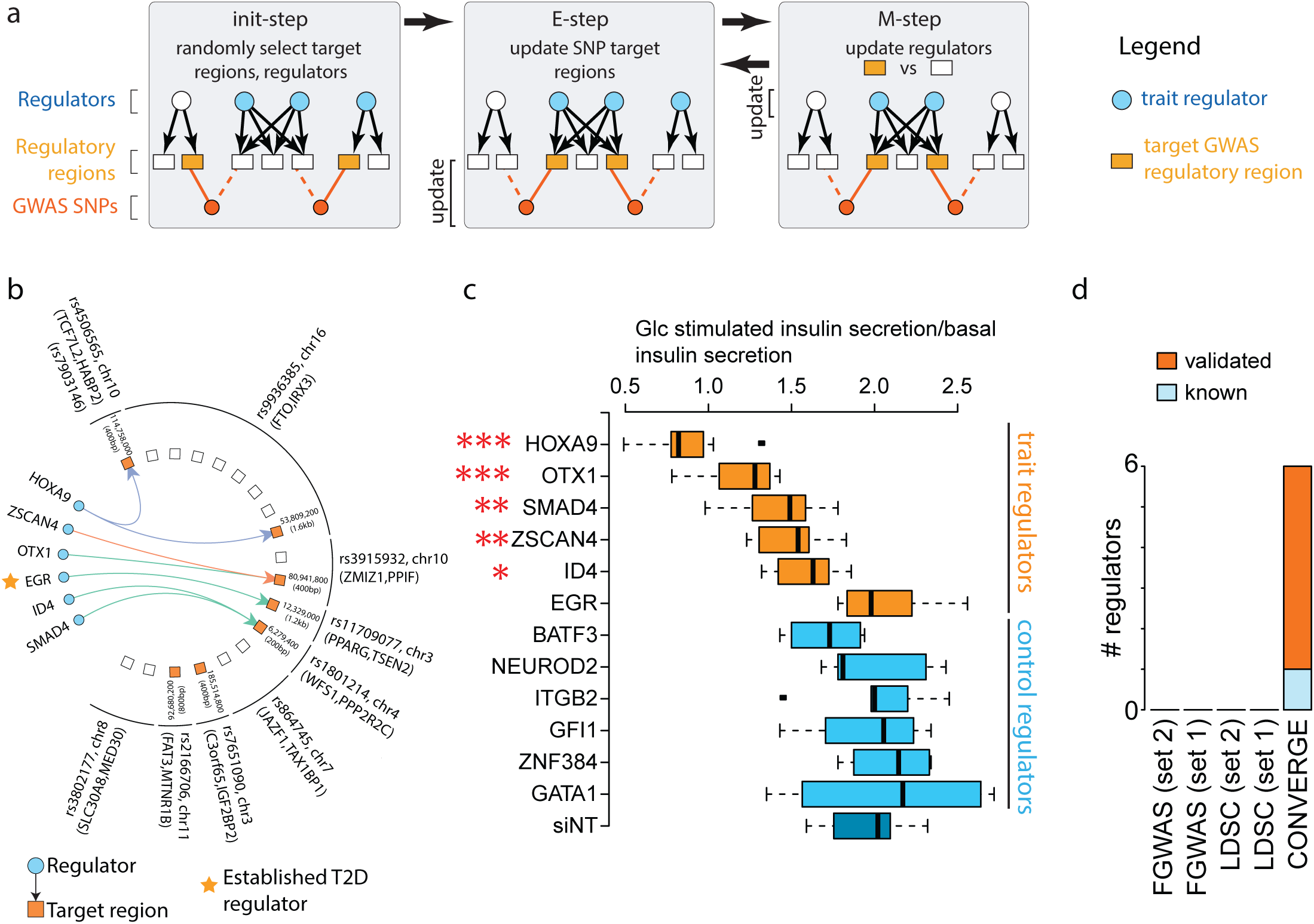
Inferring the regulatory architecture of complex traits using CONVERGE. **(a) Expectation maximization (EM) algorithm for learning CONVERGE. Initialization:** We first randomly select a set of trait regulators (blue circles) and target regulatory region for each GWAS SNP (solid orange squares). Dashed orange lines indicate regulatory regions tagged by a GWAS SNP, but not selected as the target regulatory region. **E-Step:** We update the target GWAS regulatory regions for each GWAS SNP by selecting the regulatory region within each locus that is most strongly bound to the current set of trait regulators. **M-step:** We update the selected trait regulators based on which regulators most preferentially bind (and distinguish) GWAS target regulatory regions from the remaining regulatory regions in the trait tissue. (**b**) **Network of regulators and target regulatory regions identified in pancreatic islets for type 2 diabetes.** We applied CONVERGE to identify trait regulators (blue circles) and GWAS target regulatory regions (orange squares) for type 2 diabetes in the context of the pancreatic islet network. White squares indicate regulatory regions that were not identified as trait regulatory regions. Regulators are grouped according to similarity with respect to bound regulatory regions in the pancreatic islet network (before GWAS analysis), and edges are colored based on regulator group. Target regulatory regions are labeled by their coordinates. Regulatory regions tagged by the same GWAS variant (outer arcs) are grouped together and labeled with the name of the lead GWAS variant and flanking genes. A star indicates an established T2D regulator. Note the regulatory regions not tagged by GWAS loci act as background and are also used to identify regulators but are not shown here. (**c**) **siRNA-based knockdown of predicted type 2 diabetes regulators yields T2D-consistent phenotype**. All six predicted T2D regulators were knocked down (orange boxplots), as well as six control regulators (blue boxplots) selected based on permutation testing. Loss in glucose-stimulated insulin secretion relative to unstimulated secretion is consistent with etiological hallmarks of type 2 diabetes. Stars indicate statistical significance compared to siNT (two-sided Wilcox rank sum test; one star, P < 0.05; two stars, P < 0.01; three stars, P < 0.001). (**d**) **Performance of CONVERGE, fgwas and LD Score at recovering established or validated regulators of T2D**. Columns indicate the number of predicted regulators made by each method. Fgwas and LD Score were run using two different sets of TF-SNP annotations, either those used for CONVERGE (Set 1), or those provided by another group^14^ (Set 2). Bar color indicates the number of predicted regulators that were either experimentally validated or are established regulators in the literature.

CONVERGE makes two assumptions about trait pathways:

1. Trait regulators collectively bind to trait-regulatory regions, which in turn regulate the expression of trait-genes that influence the phenotype. GWAS loci target a subset of these trait-regulatory regions.
2. Trait-regulatory regions can be distinguished from non-trait regulatory regions based on over-representation of binding sites of trait regulators.

From these two assumptions, we arrive at the following conclusions:

1. If the target regulatory regions of each GWAS locus are known, the trait regulators could be identified by finding regulators whose binding sites are collectively over-represented in the GWAS target regions, compared to other regulatory regions in the genome.
2. Similarly, if the trait-regulators were known, we can identify the GWAS target regulatory region at each GWAS locus as the one bound by the most trait-regulators, and prioritize novel regulatory regions also based on which ones are bound by trait regulators.

Because neither the trait regulators nor the GWAS target regions are known beforehand, CONVERGE uses the expectation maximization algorithm to iteratively infer both sets of unknowns (Fig. 3a). What makes this inference procedure tractable is that approximately half of the GWAS loci tag only a single regulatory region in its locus and are therefore unambiguous with respect to the target regulatory region. These unambiguous GWAS loci help narrow down the set of candidate trait regulators and guide the target region selection for the ambiguous GWAS loci (that tag multiple regulatory regions).

## Results: CONVERGE partitions ambiguous GWAS variants into cell type specific pathways

For each trait tissue of each GWAS from Figure 2a, we applied CONVERGE to identify trait-pathways, and found statistically significant trait pathways for nine combinations of GWAS and trait tissues (Supplementary Table 2), including type 2 diabetes, total cholesterol, LDL and HDL cholesterol levels. Of the 55% of GWAS loci that tag at least one regulatory region in a trait tissue, CONVERGE definitively assigns a median of 16% of GWAS loci to a trait pathway in a trait tissue. While significant trait pathways were typically only detected in one trait tissue for a GWAS, CONVERGE identified three trait-pathways for LDL cholesterol. Across the three trait tissues, 46% of variants tag regulatory regions in more than one cell type, initially suggesting they may potentially play a role in trait pathways in multiple tissue types. CONVERGE, however, assigns 42% of these ambiguous loci to one specific trait pathway in exactly one target cell type, suggesting by combining information across multiple loci we can disambiguate these loci.

## Results: CONVERGE identifies trait regulators of type 2 diabetes and total cholesterol

CONVERGE predicted a median of 50 trait regulators per study that play key roles in phenotypic variation. We began with analysis of T2D as only one trait tissue (pancreatic islets) and six corresponding trait regulators were predicted (HOXA9, ZSCAN4, OTX1, EGR, ID4 and SMAD4), allowing us to comprehensively validate all predicted regulators for this trait (Fig. 3b,c). Only EGR had prior evidence of a role in T2D, though OTX1 and HOXA9 are from the homeobox family for which other members have been implicated in type 2 diabetes pathogenesis and insulin sensitivity and secretion^42^. The prediction of HOXA9 was made because HOXA9 binding sites are present in two of the nine T2D loci in the islet network (rs4506565 at the TCF7L2 locus and rs9936385 at the FTO locus, Fig. 3b, purple lines), more than expected based on HOXA9 binding across islet regulatory regions in general (HOXA9 is in the bottom 8% of factors when ranked by promiscuity). Of the two T2D loci to which HOXA9 mapped, the GWAS locus at TCF7L2 had only one regulatory region in its locus and was therefore unambiguous with respect to the target regulatory region, which in turn helped identify HOXA2.

The causal regulatory region predicted by CONVERGE at TCF7L2 overlaps the rs7903146 variant, which has been established as the most likely causal variant due to its impact on chromatin and regulatory activity^43^.

siRNA-mediated knockdown of 5 of 6 of the predicted factors showed no effect on basal insulin secretory rate, but showed a significant reduction in glucose-stimulated insulin secretion in our assay (Fig. 3c), yielding a T2D-consistent phenotype. Strikingly, knockdown of each of the six controls selected based on permuted networks that maintain node degree yielded no such change in glucose-stimulated insulin secretion. This confirms the ability of CONVERGE to recognize trait regulators that are specific to the trait and not simply a hub in the genomic network.

To test the hypothesis that CONVERGE identifies regulator enrichments that are not the result of direct overlap of causal variants with transcription factor binding sites, we ran both fgwas^13^ and LD Score^12^ to identify globally enriched TFBS in T2D GWAS loci (Fig. 3d). We first confirmed both fgwas and LD Score were able to recover pancreatic islets as a trait tissue of T2D (Supplementary Figures 7-8), then predicted enriched TF binding sites using two separately collected sets of TF annotations: those used to construct CONVERGE’s genomic networks, and also those provided by another group^14^. We did not find any enriched TFBS globally across the T2D loci using either method or either set of annotations (Fig. 3d). We repeated our CONVERGE analysis testing multiple thresholds for determining variants in LD with the lead GWAS variants and found our T2D regulator set was highly robust (Supplementary Fig. 9). Taken together, our results suggest there exists some regulators that strongly co-localize with a subset of T2D GWAS loci, but do not directly overlap candidate causal variants.

We next turned our attention to total cholesterol (TC), as the role of the liver in its regulation is well characterized. CONVERGE predicted 65 total cholesterol regulators in adult liver (Fig. 4a), including 15 factors previously established in lipid and cholesterol homeostasis. The predicted regulators also established in the literature include SREBP1a, SREBP1c and SREBP2, three members of the sterol response family known to regulate *de novo* cholesterol synthesis and uptake of plasma lipoproteins^44^. We also recovered the nuclear receptor signaling genes FXR and RXR, which regulate cholesterol homeostasis via absorption, remodeling, transport, and synthesis and are known to convert cholesterol to bile acids^45,46^.

**Figure 4:**
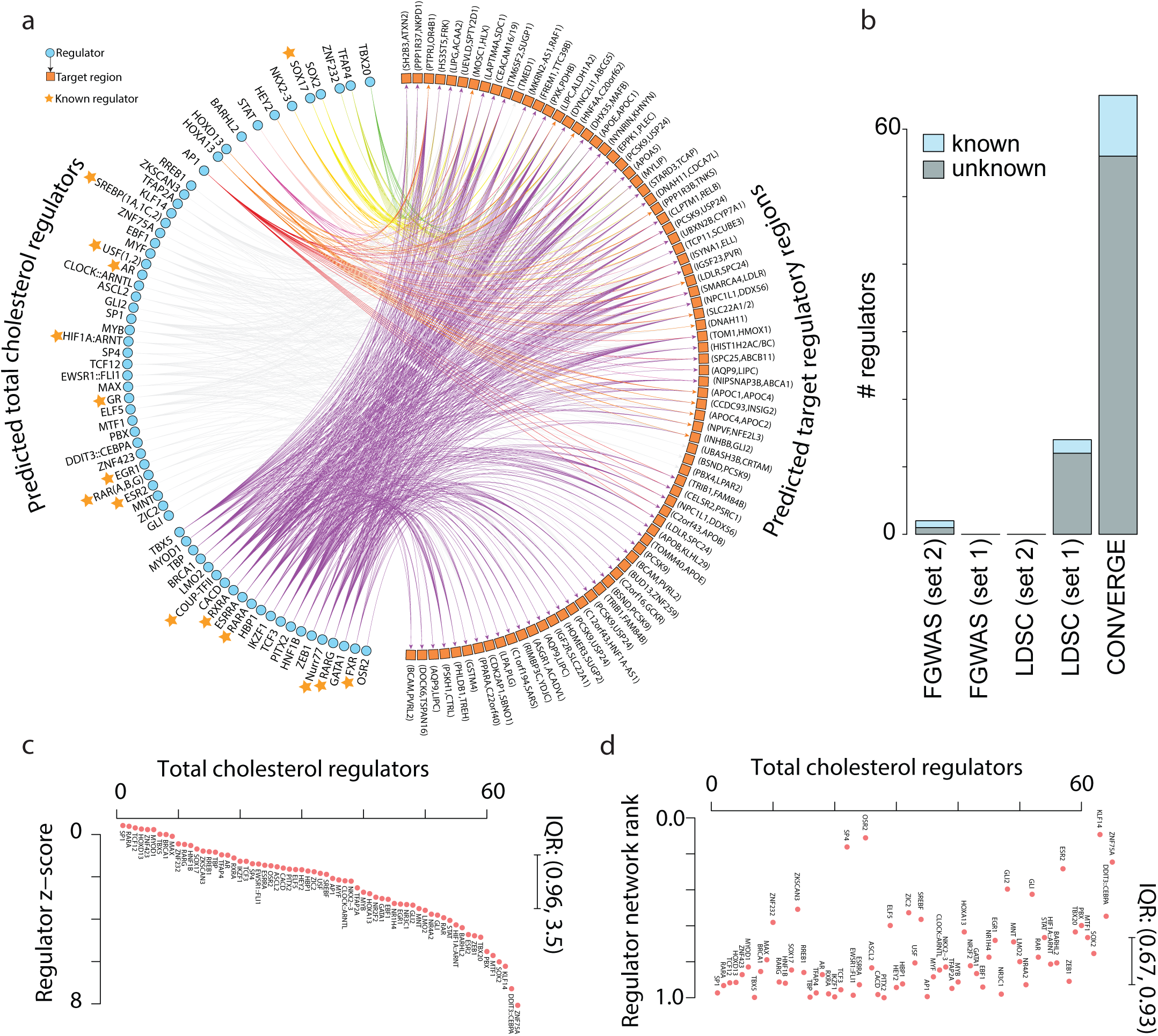
Inferring the regulatory architecture and regulator specificity of the total cholesterol pathway in liver using CONVERGE. (**a**) **The total cholesterol pathway in liver**. The total cholesterol liver network is drawn similarly to Figure 3b. (**b**) **Performance of CONVERGE, fgwas and LD Score at recovering known regulators of total cholesterol**. Columns indicate the number of predicted regulators made by each method. Fgwas and LD Score were run using two different sets of TF-SNP annotations, either those used for CONVERGE (Set 1), or those provided by another group^14^ (Set 2). Bar color indicates the number of predicted regulators that are established regulators in the literature. (**c**) **Total cholesterol regulators ordered by z-score weight.** Regulator weights *α* learned by CONVERGE using the adult liver network are scaled by the mean and standard deviation of their corresponding weights learned by CONVERGE on the 500 permuted liver networks. Higher z-scores indicate higher specificity with respect to total cholesterol regulatory regions. IQR: inter-quartile range (range of the center 50% of z-scores). (**d**) **Network rank of total cholesterol regulators.** Network rank is calculated as the relative number of times a regulator was selected as a trait regulator under the same permutation testing as in (a). Higher network rank indicates that a regulator is likely to be a hub in the network, and therefore less specific to TC. Regulators are in the same order as (a).

We compared CONVERGE to fgwas and LD Score for their ability to predict established total cholesterol regulators. Both fgwas and LD Score captured liver as the most significant trait tissue (Supplementary Figures 7-8), again confirming we successfully ran their pipelines. In terms of trait regulators, fgwas only identified POLR2A and E2F1, while LD Score predicted 14 regulators (CREB3, EHF, ELF4, ESR2, EVX2, GRHL1, RF7, KLF16, KLF7, MNT, NFYA, PTEN, SMAD4, SRF), of which two of them are established TC regulators (ESR2, GRHL1) (Fig. 4b). Both LD Score and CONVERGE recovered a similar proportion of established TC regulators (~16%) (Fig. 4b), though CONVERGE predicts more total regulators. Only ESR2 was predicted by both CONVERGE and LD Score, again demonstrating their complementary predictions. We again repeated our CONVERGE analysis testing multiple LD thresholds, and found our predicted regulators were also consistent across multiple LD thresholds, albeit with a narrower range than seen with T2D (Supplementary Fig. 10).

To characterize the novel total cholesterol regulators not yet reported in the literature, we next gauged the specificity of the total cholesterol regulators relative to global regulators of liver using two metrics: regulator z-score (measuring relative enrichment of trait regulatory regions in a regulator’s targets) and regulator network rank (relative number of times a regulator is expected to be selected as a trait regulator by chance – see Methods). TC regulators are specific to regulatory regions tagged by GWAS loci (z-score IQR is 0.96-3.5, Fig. 4c). Furthermore, while most TC regulators are not hubs of the liver network, they are important regulators in liver (network rank IQR is 0.67-0.93, Fig. 4d). Taken together, these two observations suggest that TC GWAS regulatory regions are concentrated in a liver subnetwork defined by the colocalization of trait regulators, and are not simply being connected by hub regulators in the liver network.

Surprisingly, a handful of TC regulators are characterized by high regulator z-scores but low network rank, including KLF14, GLI2, and ESR2. Variants upstream of KLF14 have been consistently associated with HDL cholesterol levels and type 2 diabetes^28,47^, and in adipose tissue, proximal variants act as a cis-eQTL to KLF14 and have a master trans-effect on genes implicated in metabolic traits^48^. KLF14 is dysregulated in the liver of dyslipidemia mice, and evidence points to KLF14 regulation of plasma HDL levels^49^. Also, therapeutic treatment of mice increased HDL levels via KLF14-mediated mechanism^49^. GLI2 is one of the flanking genes of a genome-wide significant total cholesterol SNP, despite the fact that CONVERGE did not use this information when inferring GLI2. Furthermore, the Gli family of transcriptional regulators are a key component of the Hedgehog (Hh) signaling pathway, and while cholesterol modification of Hh ligands is critical for signal transduction in Hh, recently Hh signaling has been implicated in cholesterol metabolism itself^50^. ESR2 is already an established regulator of total cholesterol, and our network uncovers its unusually high specificity to this phenotype.

## Results: Novel predicted loci harbor new GWAS variants

In addition to identifying the trait pathway, CONVERGE prioritizes the remaining regulatory regions in the genome (not tagged by GWAS variants) with respect to how strongly they are bound by GWAS trait regulators. We expect that if CONVERGE inference of trait regulators is accurate, regulatory regions highly prioritized but not proximal to GWAS variants would harbor novel GWAS variants and be proximal to new trait genes. To test this hypothesis, first, we trained CONVERGE using historical GWAS of total cholesterol, HDL and LDL cholesterol, as summary statistics of these traits were available from multiple time points, and the adult liver tissue was a statistically significant trait-tissue for all traits from all time points. We prioritized regulatory regions genome-wide using the historical data, and found we had some power to predict the locations of novel genome-wide significant loci identified in more recent studies for all three traits (Fig. 5a). This validation approach was limited by the small number of traits for which powerful GWAS were conducted both historically and recently, so we also tested whether we could predict locations of suggestive GWAS loci (5×10^−8^ < P < 5×10^−6^) found in the same study used to train CONVERGE. We found that the most highly prioritized regulatory regions identified by CONVERGE were also significantly enriched for suggestive GWAS loci for T2D, total cholesterol and LDL cholesterol (Fig. 5b). Our model therefore accurately prioritizes additional loci that are involved in the complex trait, suggesting that inference of both the trait regulators and the genomic network is accurate. Using the priority values of the suggestive GWAS loci, we conservatively identified a median of 247 novel trait regulatory regions (not tagged by any GWAS locus), which is 4.5 times as many regulatory regions as is currently tagged by genome-wide significant loci (Supplementary Fig. 11). This suggests there are many genomic locations that may still harbor un-detected variants associated with these complex traits.

**Figure 5:**
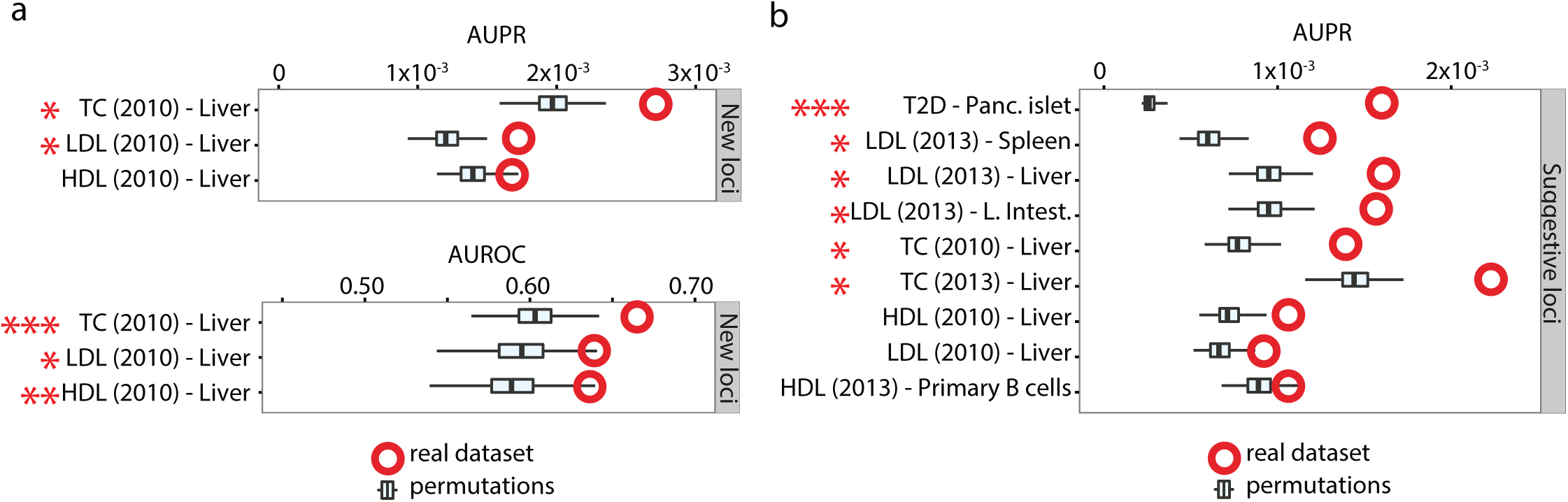
Validation of predicted novel trait regulatory regions. (**a**) **Prediction of novel GWAS regulatory regions identified in recent GWAS not used to train CONVERGE**. Both AUPR and AUROC are separate measures of prediction accuracy, where larger numbers indicate better prediction. The red open circles represent predictive accuracy using the real liver network, and the boxplots indicate the predictive performance when the liver network was permuted (see Methods). Stars indicate statistical significance compared to 500 permutations (empirical P-values; one star, P < 0.05; two stars, P < 0.01; three stars, P < 0.002, the smallest achievable). (**b**) **Prediction of suggestive GWAS regulatory regions.** For all GWAS and trait tissues for which statistically significant pathways were found, we measured CONVERGE accuracy when predicting regulatory regions tagged by suggestive GWAS loci (5×10^−8^ < P < 5×10^−6^) that were not used to train CONVERGE. We only report AUPR here because we expect only a small subset of suggestive loci to be true positive GWAS loci, an assumption better reflected by AUPR performance.

We further investigated the top prioritized novel loci for both type 2 diabetes (Fig. 6a) and total cholesterol (Fig. 6c) when CONVERGE is trained with the most recent data available. The top ranked novel locus for T2D is located in the 3q region upstream of both the STAG1 and TMEM22 transcription start sites, and is further supported by external evidence of DNase hypersensitivity (Fig. 6b). The genes in this locus have been previously implicated in diabetes nephropathy by a GWAS SNP rs1866813 (Fig. 6b)^51^ downstream of the neighboring IL20RB gene. The second most highly prioritized locus is a regulatory region located in the first intron of DR1, which is a TBP-associated repressor proposed to be under insulin control and up-regulated under loss of insulin action, and even further upregulated when diabetes is induced after loss of insulin^52^. Finally, CONVERGE identified an intergenic enhancer 4.4kb upstream of the lipoprotein lipase (LPL) gene as the fourth most prioritized region. LPL facilitates the removal of triglyceride-rich proteins from the bloodstream, and its dysregulation in diabetes contributes to the dyslipidemia of T2D^53^.

**Figure 6:**
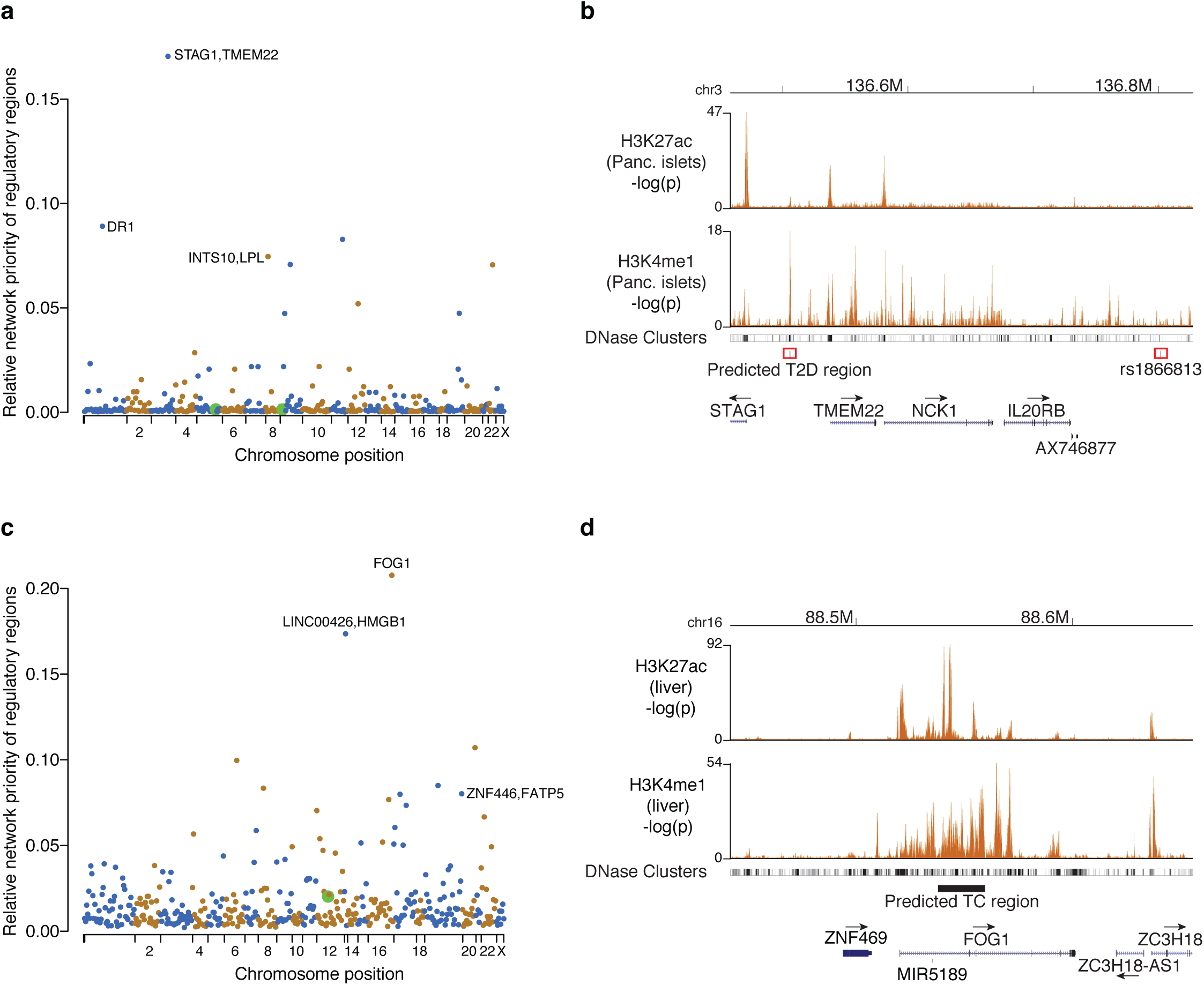
Prioritized regulatory regions are proximal to genes also implicated in similar complex traits. (**a**) **Prioritization of pancreatic islet regulatory regions using type 2 diabetes GWAS variants**. Manhattan-like plot indicates the relative prioritization for all regulatory regions in the pancreatic islet network, based on training CONVERGE using T2D GWAS variants. Green circles represent regions tagged by any suggestive GWAS SNP (5×10^−8^ < P < 5×10^−6^). Top prioritized regions are labeled with the nearest flanking genes. (**b**) **The top prioritized regulatory region for T2D is proximal to a diabetic nephropathy variant**. The top prioritized regulatory region for T2D sits upstream of the transcription start sites of a cluster of genes (STAG1, TMEM22, NCK1, IL20RB). This regulatory region is defined by the strongest H3K4me1 peak in the region, and is supported by separate DNase hypersensitivity data from a range of cell types collected by the ENCODE consortium. The regulatory region is also proximal to rs1866813, a variant associated with diabetic nephropathy. (**c**) **Prioritization of adult liver regulatory regions using total cholesterol GWAS variants**. Figure is in the same style as (a). (**d**) **The top prioritized region for total cholesterol is located in an intron of a candidate TC regulator**. This element is supported by both H3K27ac and H3K4me1 marks. FOG1 is a co-factor of GATA4, one of the regulators of lipid metabolism among other functions.

For total cholesterol, a predicted enhancer intronic to the gene FOG1 (Friend of GATA-1) was the most highly prioritized (Fig. 6d). FOG proteins are co-factors of GATA, and in the context of lipid regulation, FOG proteins as well as GATAs are implicated in adipogenesis^54^ and lipid metabolism^55^. CONVERGE also independently identified the same liver regulatory regions at the FATP5 (Fatty Acid Transport Protein 5) and HMGB1 loci, for both total cholesterol and LDL cholesterol. The primary role of FATP5 is in uptake of long-chain fatty acids, and is critical to liver lipid homeostasis^56,57^, while HMGB1 is a nuclear transcriptional regulator that plays a critical role in the liver inflammation response to elevated free cholesterol levels^58^.

## Discussion

CONVERGE is the first method to identify regulators whose binding sites co-localize with GWAS loci but do not necessarily directly overlap them. This is in contrast to GWAS fine-mapping methods that search for regulators whose binding sites directly overlap causal variants. We therefore expect CONVERGE to identify complementary sets of trait regulators. This was evident in our results, as CONVERGE was the only method able to make statistically significant regulator predictions for T2D, and there was little to no overlap of predictions between CONVERGE and other methods for total cholesterol, although the fraction of recovered established regulators were similar. We therefore envision geneticists using a combined approach involving both CONVERGE and fine-mapping methods to make mechanistic inferences.

In our type 2 diabetes experiments, all three tested methods (CONVERGE, LD Score, fgwas) were unable to recover the two factors previously reported as globally enriched in T2D GWAS loci (FOXA1/2, RFX). First, we note that FOXA1/2 is a promiscuous factor in our pancreatic islet genomic network – it is in the top 5% of regulators with respect to number of regulatory regions bound. Because CONVERGE identifies trait regulators as those that preferentially bind to GWAS loci compared to the remainder of the epigenome, such prolific TFs will be biased against by design. We reasoned that so-called hubs in the genomic network are likely to have non-specific effects on GWAS traits and are therefore of less interest. Second, the previous studies used TF ChIP-seq peaks called specifically in islet cells to identify the FOXA1/2 and RFX enrichment, while our TF-regulatory region links were defined only using in silico TFBS predictions. We did not use ChIP-seq peaks to define our regulator-regulatory region links because ChIP-seq peaks are cell type-specific, and have only been systematically collected for many TFs in a few select cell lines^59^.

CONVERGE shares some similarity with gene network prioritization methods that use the guilt-by-association principle to identify genes that interact with GWAS genes to shed light on the underlying trait pathways^60–63^. What distinguishes CONVERGE (and the fine mapping methods we compare it to here) from the gene centric network methods is that gene network-based approaches rely on pre-determined assignments of GWAS variants to target genes, even though more than 88% of GWAS variants do not tag coding variants. While these GWAS variants are enriched in regulatory regions, each GWAS variant can tag as many as 49 different regulatory regions, and approximately 60% of regulatory regions are estimated to target a non-nearest gene^64^, making identification of the causal variant (and therefore the target GWAS genes) challenging. CONVERGE avoids the task of assigning variants to genes by instead hypothesizing GWAS variants may be connected through the regulatory regions they disrupt, and thus the guilt-by-association occurs through the regulator nodes of the genomic network (as opposed to directly via gene-gene interactions). CONVERGE uses epigenomic maps to construct novel regulatory region-centric networks where nodes consist of variants, regulatory regions and TF regulators, thus generalizing the gene centric network to include the notion of non-coding regulatory regions.

Inference of the regulatory architecture of complex traits using multiple loci affords new insight on several fronts. In our study, we found 53% of regulatory GWAS variants tag regulatory regions in more than one cell type, making their mechanism ambiguous and leaving it unclear whether these variants act in multiple cell types or a single one. Our analysis reveals that for traits such as LDL cholesterol levels, we are able to assign these ambiguous GWAS variants to a single specific cell type based on regulatory similarity to GWAS variants that are unambiguously assigned to a cell type. This indicates at least several of these ambiguous GWAS variants target trait pathways within a single trait tissue. However, CONVERGE currently only assigns approximately 16% of GWAS variants to a clear trait pathway within an individual trait tissue. We expect to discover more subtle trait pathways by iteratively applying our framework after removing the previously detected trait pathway and variants before reapplication of CONVERGE.

One of the outstanding challenges of GWAS is to estimate how many loci remain to be discovered and to characterize them, and CONVERGE analysis identified two types of regions likely to harbor undiscovered variants. First, we conservatively estimated a median of 4.4 times more loci relative to the number of current genome-wide significant loci, by identifying regulatory regions distal to GWAS loci but are predicted to be strongly bound by GWAS trait regulators. Our estimate is likely a lower bound on the true number of additional loci, as our framework can only identify common trait pathways targeted by GWAS variants in the input set. Second, both coding and non-coding variants proximal to the trait regulators themselves may harbor additional rare or private GWAS variants (though CONVERGE does not explicitly look for these), as was the case for KLF14 and GLI2 for total cholesterol.

Our study focused on the analysis of two traits, type 2 diabetes and total cholesterol, for a number of practical reasons. For type 2 diabetes, the detected trait pathway is small and limited to a single trait tissue (pancreatic islets), enabling exhaustive testing of all predicted trait and control regulators. Also, both traits are well studied in the literature, providing another way to validate predictions. We note that overall, CONVERGE detects statistically significant pathways for four out of 14 GWAS studies that we tested (Fig. 2). We envision four approaches to improving CONVERGE sensitivity. First, transfer learning can be employed for traits whose trait tissues have multiple related genomic networks available. While CONVERGE currently identifies trait pathways in each genomic network separately, if multiple related genomic networks are available (e.g. for immune cells in the case of autoimmune disorders), CONVERGE can be trivially modified to identify regulators that maximize the likelihood of the GWAS data across all related genomic networks simultaneously. This will ensure the selected trait regulators are robust to small perturbations in the genomic networks, and improve power to detect trait regulators. Second, we used a stringent Bonferroni correction factor for identifying trait tissues (Fig. 2a) that treats each of the genomic networks independently, although for example immune cells are heavily overrepresented in the 127 genomic networks. Grouping these genomic networks into tissue groups and applying a more relaxed significance criterion will increase the number of trait tissues (and therefore trait pathways) identified. Third, because CONVERGE relies on identifying shared regulatory signals between GWAS loci, its ability to identify trait pathways is directly related to the number of GWAS loci identified that are regulatory in nature. As the size of the GWAS cohorts increases and the statistical models for identifying GWAS loci improves, we expect CONVERGE to gain more power to detect trait pathways in the years to come. Fourth, the 127 genomic networks do not cover all the possible trait tissues. Inference of more genomic networks using other data types as they become more available, e.g. ATAC-Seq, will allow detection of additional trait pathways.

A limitation of our approach to constructing genomic networks is the prediction of TF binding to regulatory regions via combining position weight matrix (PWM) scanning with epigenomic annotations, instead of directly using cell type specific TF ChIP-seq datasets, which are comparatively rare but more accurate than PWM scanning. Multiple TFs, particularly from the same family, may share similar binding preferences, and therefore some regulator nodes are not strictly identifiable. First, despite this limitation, we have shown via systematic knockdown of all predicted regulators as well as controls, that CONVERGE is still able to identify bona fide T2D regulators. Second, DNA binding domains with similar protein sequence also tend to share DNA sequence preferences, providing a means for deconvoluting which set of TFs is represented by each ambiguous regulator node, that can then further be deconvoluted if TF expression data is available^65^. Third, the prioritization of regulatory regions (both within GWAS loci and outside) does not depend on the identity of the regulator nodes – for example, regulator nodes could also represent de novo motifs that have not been matched to known TFs but nonetheless are useful for connecting regulatory regions bound by a common unknown TF. Finally, regulator-regulatory region connections in the genomic networks can be refined using e.g. ChIP-Seq peaks of known transcription factors such as those used to validate our genomic networks.

## Description of Supplemental Data

Supplemental Data includes 15 supplementary figures, two supplementary tables, and supplementary methods.

## Declaration of Interests

The authors declare no competing interests.

## Web Resources

### Code availability

All data and code will be released on GitHub (https://github.com/gquon) upon publication, and can be found online currently at http://people.csail.mit.edu/geraldquon/CONVERGE_all.zip.

### Data availability

Publicly available summary statistics used in this study are available here: https://ucdavis.box.com/s/7s9wldqk7iki7hhlilaas5f8e9qol464, and will be released on GitHub as well upon publication.

